# Submaximal eccentric resistance training increases serial sarcomere number and improves dynamic muscle performance in old rats

**DOI:** 10.1101/2024.07.10.602916

**Authors:** Avery Hinks, Ethan Vlemmix, Geoffrey A. Power

**Author notes:** **Correspondence:** Geoffrey A. Power PhD., **N**euromechanical **P**erformance **R**esearch **L**aboratory, Department of Human Health and Nutritional Sciences, College of Biological Sciences, University of Guelph, Ontario, Canada.

## Abstract

The age-related loss of muscle mass is partly accounted for by the loss of sarcomeres in series. Serial sarcomere number (SSN) influences muscle mechanical function including the force-length and force-velocity-power relationships, and the age-related loss of SSN contributes to declining performance. Resistance training biased to active lengthening (eccentric) contractions increases SSN in young muscle, however, we showed maximal eccentric training in old rats did not alter SSN and further worsened performance. A submaximal eccentric training stimulus may be more conducive to adaptation for aged muscle. Therefore, the purpose of this study was to assess whether submaximal eccentric training can increase SSN and improve mechanical function in old rats. Twelve 32-month-old male F344/BN rats completed 4 weeks of submaximal (60% maximum) isokinetic eccentric plantar-flexion training 3 days/week. Pre- and post-training, we assessed *in-vivo* maximum isometric torque at a stretched and neutral ankle angle, the passive torque-angle relationship, and the isotonic torque-angular velocity-power relationship. The soleus and MG were harvested for SSN measurements via laser diffraction, with the untrained leg as a control. SSN increased 11% and 8%, and muscle wet weight increased 14% and 13% in the soleus and MG, respectively. Maximum isometric torque and shortening velocity were unaltered, but there was a shift towards longer muscle lengths for the optimal angle of torque production, a 42% reduction of passive torque, and 23% increase in peak isotonic power. Eccentric training at 60% maximum was beneficial for aged muscle, increasing SSN and muscle mass, reducing muscle passive tension, and improving dynamic performance.

**New & Noteworthy:**

- Four weeks of submaximal (60% of maximum) eccentric resistance training increased serial sarcomere number and muscle mass in old rats
- Increased serial sarcomere number corresponded to broadening of the torque-angle relationship following training, which contributed to increased peak isotonic power
- Submaximal eccentric training also led to a large reduction in passive torque throughout the range of motion

## Introduction

Muscle fascicle length (FL) becomes shorter with age driven by a loss of in-series aligned sarcomeres (1–4). This loss of serial sarcomere number (SSN) contributes to muscle atrophy and impaired mechanical function in old age (3, 5). We recently showed that aged muscle retains the ability to re-add sarcomeres in series following disuse atrophy, and SSN was notably more adaptable than measures of parallel muscle morphology (i.e., muscle thickness and pennation angle) (6). Hence, increasing FL by adding sarcomeres in series represents an appealing target for training interventions to improve performance in older adults.

The most common exercise intervention to stimulate sarcomerogenesis is resistance training biased to active lengthening (eccentric) contractions (7, 8). In young healthy rats, eccentric training can increase SSN by up to 8% (4, 9–11), and in humans SSN adaptations have been assumed via increases in FL as measured by ultrasound (12, 13). Increases in FL following submaximal (60-80% maximum) eccentric training have also been reported in older adults (14–16), however, assuming sarcomerogenesis from ultrasound-derived FL holds limitations without simultaneous measurement of sarcomere length (SL) (17). Direct measurement of SL in humans is invasive with limited accessibility (18), therefore, animal models are needed to investigate the ability for eccentric training to induce sarcomerogenesis in aged muscle.

We recently showed maximal eccentric training induced sarcomerogenesis and beneficial mechanical adaptations in young rats, however, sarcomerogenesis was blunted and function was worsened in old rats (4), likely due to an age-related impaired recovery from muscle damage following high-intensity eccentric exercise, leading to dysfunctional remodeling (19). Therefore, the purpose here was to investigate if aged muscle can add sarcomeres in series and improve mechanical function following submaximal eccentric training.

## Methods

### Animals

Twelve 32-month-old male Fisher 344/Brown Norway rats were obtained (Charles River Laboratories, QC, Canada). The University of Guelph’s Animal Care Committee (AUP #4905) approved all protocols. Rats were housed at 23°C in groups of two or three and given ad-libitum access to a Teklad global 18% protein rodent diet (Envigo, Huntington, Cambs., UK) and room-temperature water. Pre-training mechanical testing was completed at 32 months of age then training commenced 4-6 days later. Training lasted 4 weeks, then post-training mechanical testing was completed 72 hours following the final training session. The soleus and MG were dissected for SSN determinations. The left leg completed training while the right leg acted as an internal control in accordance with previous studies (4, 17, 20, 21).

### Data acquisition during mechanical testing and training

A 701C High-Powered, Bi-Phase Stimulator (Aurora Scientific, ON, Canada) was used to evoke transcutaneous muscle stimulation as described previously (4). Torque, angle, and stimulus trigger data were sampled at 1000 Hz with a 605A Dynamic Muscle Data Acquisition and Analysis System (Aurora Scientific, ON, Canada).

### Mechanical testing

As described previously (4), rats were anesthetized with isoflurane and positioned supine on a heated platform (37°C). The left leg was shaved and fixed to a force transducer/length controller foot pedal via tape, with the knee immobilized at 90°. Each mechanical testing session began with determination of the optimal stimulation current for active plantar flexion torque (frequency = 100Hz, pulse duration = 0.1ms, train duration = 500ms) at an ankle angle of 90° (full plantar flexion = 180°), which was the current used throughout the remainder of the testing session. This stimulation current was confirmed to maximally activate the plantar flexors with minimal spread to the antagonist as described previously (4). A 100Hz stimulation was then completed at an ankle angle of 70°. Active torque was measured by subtracting the minimum value at baseline (passive torque) from the total torque during stimulation (11, 22). A passive torque-angle relationship was then constructed by recording the minimum passive torque following 5s of stress-relaxation at ankle angles of 100, 95, 90, 85, 80, 75, and 70°. A torque-angular velocity-power relationship was then constructed from isotonic contractions. For each isotonic contraction, the ankle started at 70°, then contracted against a load clamp to a maximum angle of 110°. Isotonic (i.e., constant torque) contractions were performed at load clamps equating to 10, 20, 30, 40, 50, 60, 70, and 80% of the maximum isometric torque at 70°, in a randomized order. Angular velocity was recorded as the maximum time derivative of the angular displacement during the isotonic contraction (4). Two minutes of rest separated each stimulation to minimize the development of muscle fatigue.

The estimated maximum shortening velocity at zero load (V_max_), maximum power (torque multiplied by angular velocity), and torque and velocity at peak power were determined by fitting the measured torque and angular velocity values to a rectangular hyperbolic curve (23):

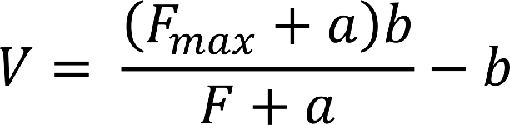

With F_max_ being maximum isometric torque at 70°, *F* and *V* representing the isotonic load clamp’s torque and angular velocity, respectively, and *a* and *b* representing Hill’s thermodynamic constants with the units of torque (N•m) and velocity (°/s), respectively. All curve-fitting was performed in Python 3 using least square error optimization.

### Isokinetic eccentric training

The submaximal eccentric training protocol was modified from our previous maximal protocol (4). Training lasted 4 weeks and occurred 3 days/week (Monday, Wednesday, Friday). Throughout training, we used a stimulation current that corresponded to 60% of the maximum isometric torque.

At the start of each session, the stimulus current was first adjusted to produce maximum isometric torque (pulse duration = 0.1ms, frequency = 100Hz, train duration = 500ms) at an ankle angle of 90°. The current was then adjusted to produce an isometric torque that as closely as possible matched 60% of maximum isometric torque. Maximum isometric torque was recorded for each training session to track changes in muscle strength throughout the training period.

Each eccentric repetition consisted of 3 phases: 1) a 500-ms pre-activation at 110°; 2) active lengthening to 70°; and 3) 3 seconds of deactivation followed by a return to 110° at 20°/s. An additional 3 seconds of rest were provided before the next repetition. To progress the eccentric training stimulus throughout, we increased the number of repetitions and the velocity of the eccentric contractions, in accordance with our previous training study (4). During week 1, rats completed 3 sets of 8 repetitions at 40°/s. In week 2, they completed 3 sets of 9 repetitions at 40°/s. Weeks 3 and 4 consisted of 3 sets of 9 and 10 repetitions, respectively, at 80°/s. Two minutes of rest were provided between each set.

### Serial sarcomere number determinations

Following post-training mechanical testing, rats were euthanized via isoflurane anesthetization followed by CO_2_ asphyxiation and cervical dislocation. The hindlimbs were amputated and fixed in 10% phosphate-buffered formalin with the ankle and knee pinned at 90°. After fixation for 1-2 weeks, the muscles were dissected and rinsed with phosphate-buffered saline. The muscles were then digested in 30% nitric acid for 6-8 hours to remove connective tissue and allow for individual muscle fascicles to be teased out (10, 11).

For each muscle, two fascicles were obtained from each of the proximal, middle, and distal regions and averaged for the reporting of data. Dissected fascicles were placed on glass microslides then FLs were measured using ImageJ (version 1.53k, National Institutes of Health, USA) from pictures captured by a level, tripod-mounted digital camera, calibrated to a ruler in plane with the fascicles. Sarcomere length measurements were taken at six locations proximal to distal along each fascicle via laser diffraction (Coherent, Santa Clara, CA, USA) with a 5-mW diode laser (∼1 mm beam diameter, 635 nm wavelength) and custom LabVIEW program (Version 2011, National Instruments, Austin, TX, USA) (24). For each fascicle, the six SL measurements were averaged to determine average SL. SSN of each fascicle was calculated as:

[uamth2]

### Statistical analysis

In SPSS Statistics Premium 28, normality of data was confirmed using Shapiro-Wilk tests. Differences in SSN, SL, FL, and muscle wet weight between the trained and untrained leg were assessed by one-way analysis of variance (ANOVA). Changes in V_max_, peak power, and torque and velocity at peak power from pre- to post-training were also assessed via one-way ANOVA. Two-way ANOVAs assessed training-induced changes in maximum isometric torque (training [pre, post] × angle [90°, 70°]), passive torque (training [pre, post] × angle [100°-70°]) and angular shortening velocity (training [pre, post] × load [10%-80%]). A Greenhouse-Geisser Correction for sphericity was applied for all ANOVAs. Significance was set at α = 0.05.

## Results

### Data exclusion

One rat died between the penultimate and final training sessions and was excluded from all analyses (now n=11). One additional rat was excluded from analyses of mechanical data due to technical issues with its experimental setup at post-training testing. Thus, the final sample size was n=11 for morphological data and n=10 for mechanical data.

### Training-induced changes in muscle morphology

Muscle wet weight increased 14% (*P*<0.001) and 13% (*P*=0.001) from the untrained to trained leg for the soleus and MG, respectively (Figure 1A-B).

**Figure 1:**
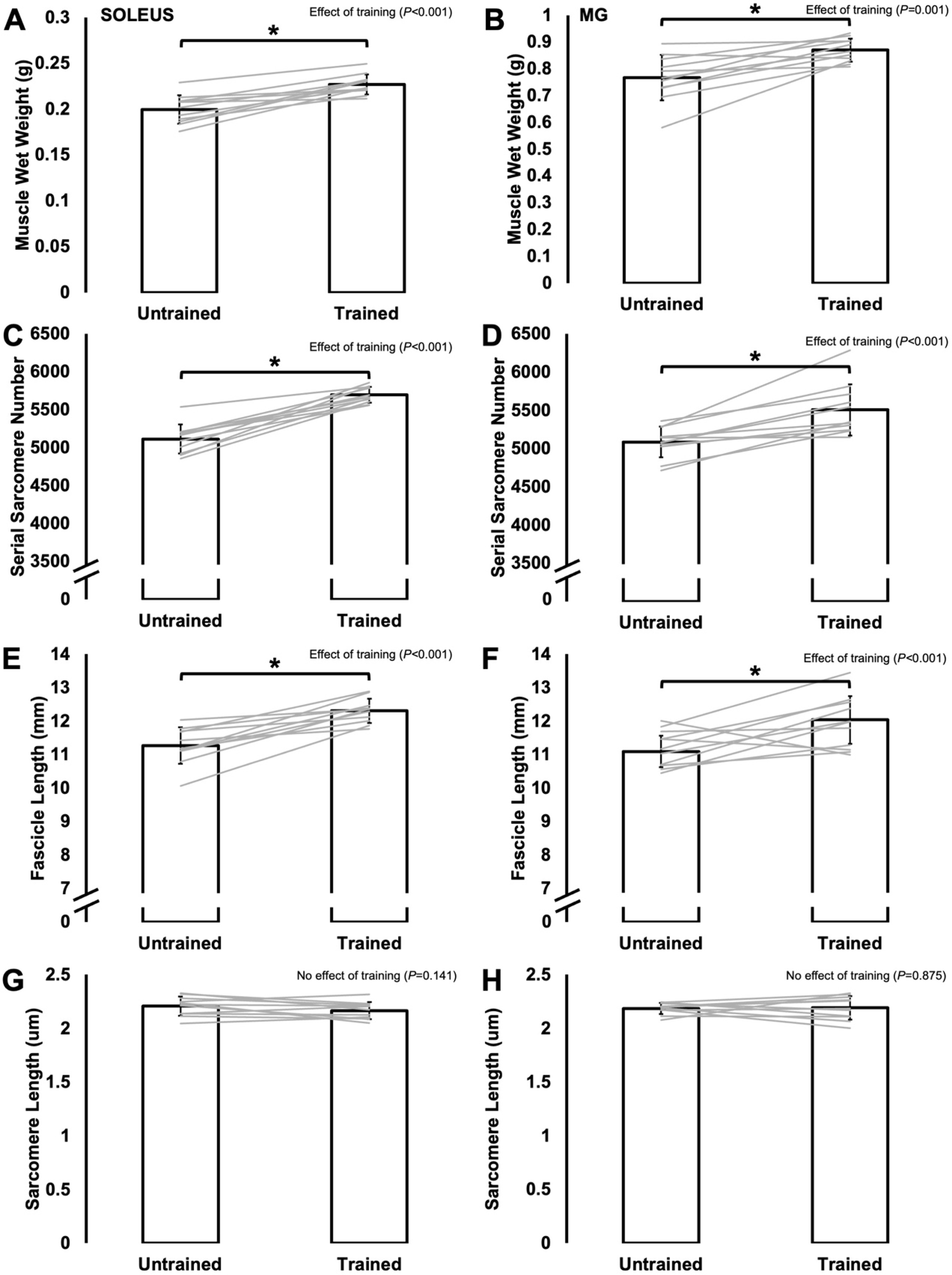
Differences in muscle wet weight (A-B), serial sarcomere number (C-D), fascicle length (E-F), and sarcomere length (G-H) between the untrained and trained leg (n=11). *Difference between trained and untrained (*P*<0.05).

SSN also increased 11% in the soleus (*P*<0.001) and 8% in the MG (*P*<0.001) from the untrained to trained leg (Figure 1C-D). SL as measured at 90° did not differ between legs (soleus: *P*=0.141; MG: *P*=0.875) (Figure 1E-F), therefore these increases in SSN resulted in 9% longer FLs as measured at 90° in both muscles (soleus: *P*<0.001; MG: *P*<0.001) (Figure 1G-H).

### Training-induced changes in muscle mechanical performance

For maximum isometric torque, there was a training×angle interaction (*P*=0.003). Neither torque at 90° (*P*=0.448) nor 70° (*P*=0.178) changed from pre- to post-training (Figure 2A). However, pre-training, torque at 90° was greater than at 70° (*P*<0.001), whereas post-training torque no longer differed between these angles (*P*=0.126) with torque at 70° being on average greater (Figure 2A). This lack of difference in torque between a neutral and stretched angle post-training indicates a training-induced shift in the torque-angle relationship such that a broader range of angles were close to optimal torque production throughout the joint range of motion.

**Figure 2:**
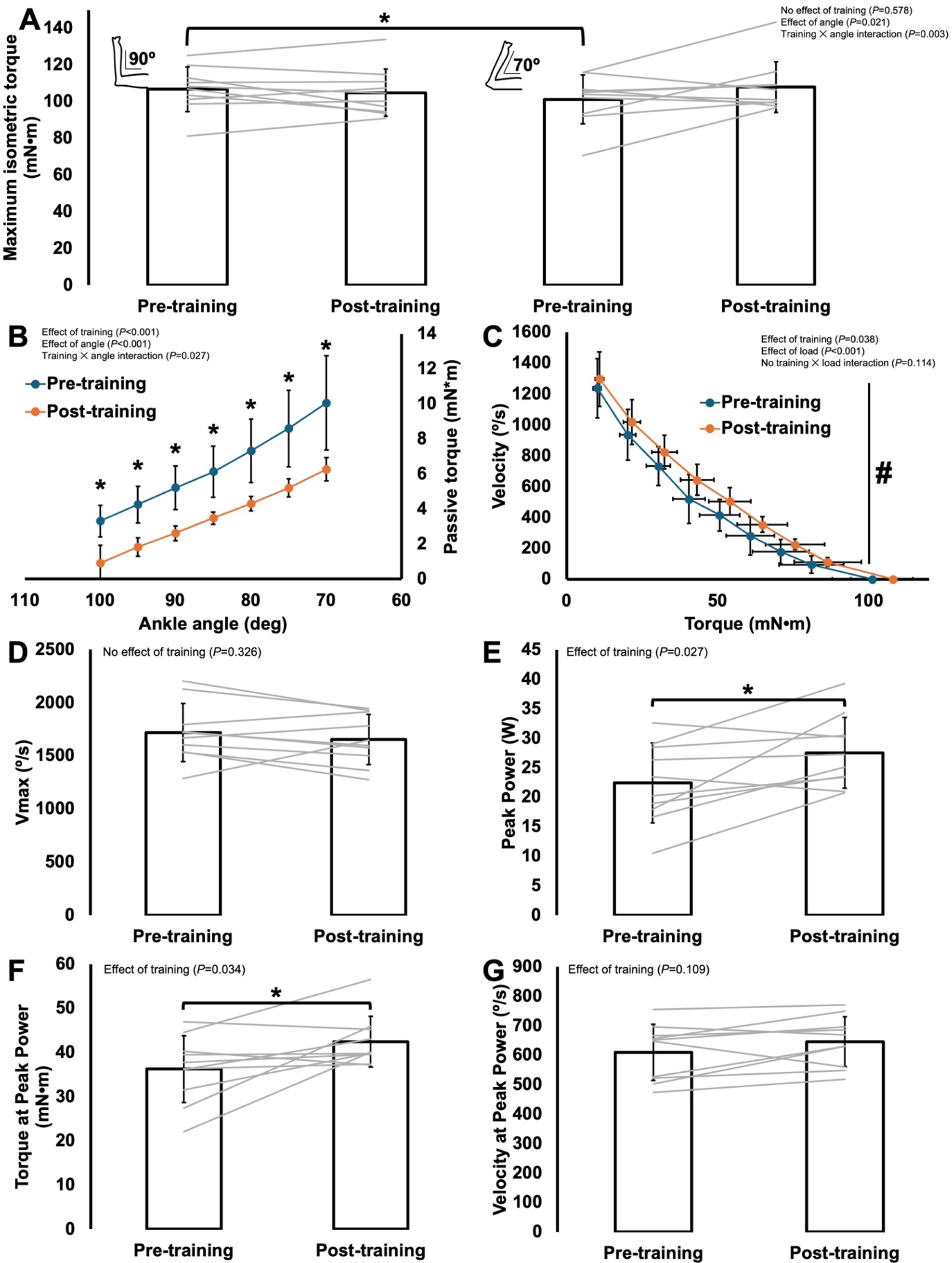
Training-induced changes in maximum isometric torque at ankle angles of 90° and 70° (A), the passive torque-angle relationship (B), the torque-angular velocity relationship (C), maximum shortening velocity (V_max_) (D), peak power (E), torque at peak power (F), and velocity at peak power (G) (n=10). A, E, and F: *Difference between points. B: *Difference between pre and post (*P*<0.05). C: #Effect of training across loads (*P*<0.05).

Torque at 90° also did not change throughout the training period (no effect of training day, *P*=0.156) unlike our previous maximum eccentric training study in old rats, in which torque at 90° progressively decreased (4) (Figure 3).

**Figure 3:**
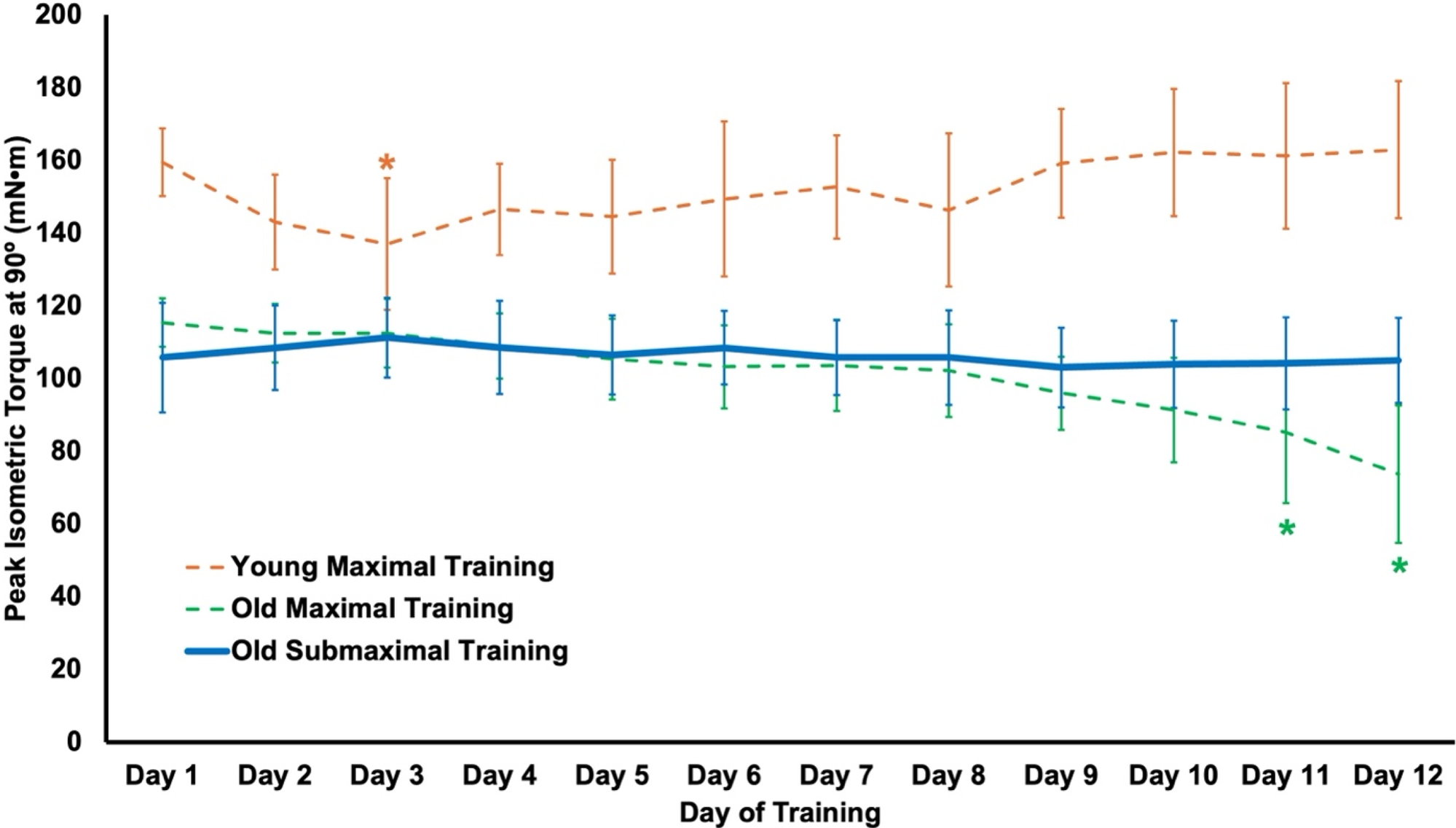
Changes in maximum isometric torque at an ankle angle of 90° throughout the training period in the present study (solid line, old submaximal eccentric training; n=10) plotted alongside our previous study’s data (4) for maximal eccentric training (dashed lines) in young (n=10) and old rats (n=11). *Difference from baseline (*P*<0.05).

For passive torque, there was a training×angle interaction (*P*=0.027). Both pre- and post-training, passive torque increased continuously as the muscles stretched from an ankle angle of 100° to 70° (all comparisons *P*<0.001). More importantly, there was a training-induced 38% to 72% reduction (all *P*<0.001) in passive torque across all joint angles (Figure 2B).

For angular shortening velocity, there was the expected effect of load such that velocity decreased with increasing load (*P*<0.001). There was also an effect of training (*P*=0.038) but no interaction (*P*=0.114), with angular velocity increasing 13% across all loads combined from pre- to post-training (Figure 2C). However, training did not affect V_max_ (*P*=0.326) (Figure 2D).

Peak power also showed an effect of training, increasing 23% (*P*=0.027) (Figure 2E). This training-induced increase in peak power was driven by increased torque rather than velocity, as torque at peak power increased 17% (*P*=0.034) (Figure 2F) while velocity at peak power did not change (*P*=0.109) (Figure 2G).

## Discussion

Previously we showed maximal eccentric training induced serial sarcomerogenesis in young rats, but in old rats did not change SSN and further worsened muscle mechanical function (4). Since eccentric contraction-induced muscle damage is more prevalent with increasing contraction intensity (25), and old rats take longer to recover from damage (19), sarcomerogenesis was likely blunted due to accumulation of muscle damage and insufficient recovery between each training session. The present study aimed to determine whether a submaximal (60% of maximum torque) eccentric training stimulus would better increase SSN and improve mechanical function in old rats, as suggested by studies on humans using loads equivalent to 50-80% of 1-repetition maximum (14–16, 26).

Indeed, we observed an 11% and 8% increase in SSN of the soleus and MG, respectively, following 4 weeks of submaximal isokinetic eccentric training. These increases in SSN manifested as 9% longer FLs, as resting SLs were unchanged (Figure 1). The increased SSN likely also contributed to the 13-14% greater muscle wet weights. While we did not account for potential addition of sarcomeres in parallel (increased fibre cross-sectional area), given the unchanged maximum isometric torque, addition of sarcomeres in parallel was likely minimal. Our findings are consistent with previous studies that observed increased SSN following eccentric training in young rats (4, 9–11, 22). Importantly, we now show that training-induced sarcomerogenesis is also possible in old rats.

In contrast to our previous maximal eccentric training study (4), we did not observe a progressive decrease in maximum isometric torque throughout the submaximal training period, and instead observed no change in torque (Figure 3). This comparison further indicates that maximal eccentric training (4) induced dysfunctional remodelling leading to impaired muscle mechanical performance, while submaximal eccentric training was a low enough intensity to allow sufficient recovery and adaptation between sessions.

We did not observe any training-induced increases in maximum isometric torque or shortening velocity. However, and perhaps more insightful into the unique adaptations to longitudinal muscle growth, we observed a broadening of the torque-angle relationship such that torque was closer to optimal throughout the joint range of motion, as indicated by torque being greater at an ankle angle of 90° (neutral) than 70° (stretched) pre-training, but no longer different between these angles post-training. A rightward-shifted, broadened force-length curve has been observed alongside increased SSN previously (11, 20, 27), and is likely associated with a reestablishment of optimal SL at a longer muscle length due to increased SSN.

Broadening of the torque-angle relationship likely also contributed to the training-induced 23% increase in peak isotonic power (Figure 2E) by making torque closer to optimal throughout the joint range of motion (28) and increasing the force component of power. This effect on the force component of power seems especially likely given torque at peak power increased 17% (Figure 2F). As angular velocity at peak power remained unchanged (Figure 2G), the velocity component of power contributed less to the improved peak power despite a modest effect of training on angular shortening velocity across all loads (Figure 2C).

Submaximal eccentric training also induced a 38-72% reduction in passive torque (Figure 3B). Decreased muscle passive force (11) and joint passive torque (29) have been observed alongside increased SSN previously, and could be due shorter SLs at a given absolute muscle length, reducing passive force contributed by sarcomeric structures (e.g., titin) (8). The reduction in passive torque could have also resulted from remodelling of the extracellular matrix, which can occur following eccentric exercise (30, 31), but those measures were beyond the scope of this study. Considering resting SL at an ankle angle of 90° did not differ between untrained and trained muscles, extracellular matrix remodelling likely accounted for most of the observed reduction in passive torque.

## Conclusion

We found that eccentric training at 60% maximum induced sarcomerogenesis in the old rat plantar flexors, suggesting that “less is more” when designing training programs for longitudinal growth of aged muscle. Concomitantly, there were improvements in muscle mechanical function including reduced passive tension and improved peak isotonic power. The increase in power is especially notable given it is a measure of dynamic muscle performance, applicable to real-world movement, and a strong predictor of function in older adults.

## Acknowledgements

This project was supported by the Natural Sciences and Engineering Research Council of Canada (NSERC). The animals were obtained from the National Institute on Aging (NIA) aged rodent colonies.

## Ethics statement

Approval was given by the University of Guelph’s Animal Care Committee and all protocols followed CCAC guidelines (AUP #4905).

## Data availability

All data generated or analysed during the study are available from the corresponding author upon request.

## Author contributions

A.H. and G.A.P. conceived and designed research; A.H. and E.V. performed experiments; A.H. analyzed data; A.H., E.V., and G.A.P. interpreted results of experiments; A.H. prepared figures; A.H. and G.A.P. drafted manuscript; A.H., E.V., and G.A.P. edited and revised manuscript; A.H., E.V., and G.A.P. approved final version of manuscript.

